# A metabolomic view on local climate adaptation: Latitudinal divergence of heat and drought responses in a coastal plant

**DOI:** 10.1101/2022.10.14.512208

**Authors:** Karin Schrieber, Svea Gluesing, Lisa Peters, Beke Eichert, Merle Althoff, Karin Schwarz, Alexandra Erfmeier, Tobias Demetrowitsch

**Affiliations:** Kiel University, Faculty of Mathematics and Natural Sciences, Institute for Ecosystem Research, Divsion Geobotany; Kiel University, Faculty of Agricultural and Nutritional Sciences, Institute for Human Nutrition and Food Science, Divsion Food Technology; Anhalt University of Applied Sciences, Department of Agriculture, Ecotrophology and Landscape Development; University of Hamburg, Institute of Plant Science and Microbiology

**Keywords:** alkaloids, *Cakile maritima*, drought, FT-ICR-MS, glucosinolates, heat, local adaptation, metabolomics, secondary metabolites, stress resistance

## Abstract

Studying natural variation in multi-stress resistance is central for predicting and managing the population dynamics of wild plant species under rapid global change. Yet, it remains a challenging goal in this field to integrate knowledge on the complex biochemical underpinnings for the targeted ‘non-model’ species. Here, we studied latitudinal divergence in combined drought and heat stress resistance in European populations of the dune plant *Cakile maritima*, by combining comprehensive plant phenotyping with metabolic profiling via FT-ICR-MS and UPLC-TQ-MS/MS.

We observed pronounced constitutive divergence in growth phenology, leaf functional traits and defence chemistry (glucosinolates, alkaloids) among population origins. Most importantly, the magnitude of growth reduction under stress was partly weaker in southern plants and associated with divergence in plastic growth responses (root expansion, leaf abscission) and the stress-induced modulation of primary and specialized metabolites with known central functions not only in plant abiotic but also biotic stress resistance.

Our study supports that divergent selection has shaped the constitutive and stress-induced expression of numerous morphological and biochemical functional traits to mediate higher abiotic stress resistance in southern *Cakile* populations, and highlights that metabolomics is a powerful tool to explore the mechanistic underpinnings of local stress adaptation in ‘non-model’ species.

**Highlight:** Plant defence chemistry and its modulation by abiotic stress exhibits latitudinal clines across natural populations of a coastal plant.

## 1 Introduction

Global change alters the composition of the stress matrix for an ever-increasing number of wild plant species (Sage, 2020; Zandalinas et al., 2021a). How species respond to such change on evolutionary time scales will dictate the likelihood of their persistence and the degree to which their natural distribution ranges may shift (Anderson and Song, 2020). The study of natural variation in plant multi-stress resistance, i.e., in the ability to maintain fitness in stressful environments, is central for predicting the future dynamics of wild plant populations and developing efficient management practice (Benito Garzón et al., 2019; Gougherty et al., 2021). Integrating knowledge on the complex biochemical underpinnings of plant multi-stress resistance for the targeted ‘non-model’ species remains a challenging, yet desirable goal in this field (VanWallendael et al., 2019), which we approach in this study.

The natural distribution ranges of many plant species span steep gradients in the frequency, intensity and quality of multiple environmental stressors. Their populations thus typically evolve divergence in stress resistance to obtain advantage in their local habitats, i.e., local adaptation (VanWallendael et al., 2019; Lortie and Hierro, 2022). Such natural variation in stress resistance provides the raw material for future adaptation to rapidly changing environments and its magnitude is thus proportional to a species’ evolutionary potential (Aubin et al., 2016). Studying natural variation in multi-stress resistance can identify populations, which are pre-adapted to future environmental change and this information may underpin species distribution modelling and conservation practice in terms of assisted gene-flow or migration (Benito Garzón et al., 2019; Bucharova et al., 2019; George et al., 2020; Gougherty et al., 2021). Moreover, the correlation structure of natural trait variation may uncover constraints to the evolution of stress resistance in the future. As resources are limited, the ability to resist one stressor can trade-off against the ability to resist another one or impose a fitness cost (Walters and Heil, 2007), which may counter adaptation to altered stress regimes.

Reciprocal transplant experiments with ecologically relevant ‘non-model’ plant species yielded broad evidence for natural variation, trade-offs and costs for multi-stress resistance (Bemmels and Anderson, 2019; George et al., 2020; Johnson et al., 2022) but left its mechanistic underpinning uncovered in large parts. Stress fosters the evolution of complex constitutive and inducible adjustments in protein and metabolite synthesis, which are complemented by diverse morphological and phenological modifications (Bhoopander and Sharma, 2020). These responses mediate the ability to maintain plant fitness (or resist stress) while either preventing or allowing that the cell is at equilibrium with the environment, i.e., avoidance or tolerance. A plants’ stress resistance is determined by a combination of many avoidance and tolerance traits, whereby the latter mainly operate at the level of the metabolome and proteome (Bhoopander and Sharma, 2020). The relative contribution of tolerance to plant stress resistance was reported to increase with stress intensity or frequency in wild populations of *Arabidopsis thaliana* (Hoermiller et al., 2018), suggesting that such strategies could be particularly relevant for species facing rapid global change. Biochemical stress responses must thus be included in the study of natural variation in the concerned species, which so far largely focussed on morpho-phenological traits with limited informative value for plant tolerance strategies (VanWallendael et al., 2019; Walker et al., 2022). Metabolomics are a promising tool to approach this aim, as they can yield insight into all states of the global stress effect (i.e., disruption of homoeostasis, stress signalling, recovery and maintenance of homoeostasis) without broad previous knowledge on the system (Villate et al., 2021). The few studies addressing metabolic stress responses in the context of natural variation have mainly focussed on herbivory or other individual stressors (Jandová et al., 2015; Calf et al., 2018; Schrieber et al., 2018, 2019; Buckley et al., 2019; López-Goldar et al., 2019). The necessity of further extending these integrative approaches to stress combinations for satisfying the complexity of natural environments and the changes they face is obvious.

Different stressors can elicit opposing plant responses, which may result in inhibitory interactions when applied simultaneously. Combined stressors thus require unique responses that cannot be extrapolated from those to individually applied stress (Zhang and Sonnewald, 2017; Zandalinas et al., 2021b). Combined heat and drought are particularly relevant in this context given their common co-occurrence across climate zones and the regional increase in their frequency and intensity predicted under global change (ICCP 2021). Heat stressed plants are known to open their stomata for reducing leaf temperature *via* transpiration, only in the absence of drought stress, which would otherwise result in a drastic reduction of leaf water potential (Rizhsky et al., 2004). Georgii *et al*. (2017) demonstrated that specific molecular response pathways, which are enhanced by combined drought and heat (e.g., growth inhibition) show no overlap with those enhanced by individually applied stressors, while other responses (e.g., maintenance of ROS homeostasis, immune responses) are enhanced under individual drought or heat but silenced or non-additively induced under combined stress. Although recent research shed light on plant responses to combined drought and heat for *Arabidopsis thaliana* and various crop plants, we still lack insight in the context of natural variation in ecologically relevant ‘non-model’ species (Nagler et al., 2018).

Here, we study latitudinal divergence in resistance to combined drought and heat in *Cakile maritima* Scop. (Brassicaceae) with an integrated approach. Our study species is native to marine coastal ecosystems in Europe where it provides a broad variety of ecosystem services, and it is widely recognised as a promising future crop plant for saline agriculture (Arbelet-Bonnin *et al*. 2019). We exposed populations originating from Northern and Southern Europe to short-term drought and heat in a fully factorial experimental design for comprehensive plant phenotyping and metabolic profiling. We hypothesized that **i)** northern and southern plants exhibit constitutive differences in growth and morphological as well as biochemical functional traits; **ii.a)** stress reduces growth in plants from both origins, whereby combined heat and drought exceed the additive effects of individually applied stressors; **ii.b)** stress induces targeted responses in morphological and biochemical functional traits in plants from both origins, whereby responses to individual heat and drought largely oppose each other, which requires a set of unique responses to the stress combination; **iii.a)** the growth reduction following stress is higher in northern than southern plants, i.e., plants from northern populations are less resistant to both stressors and **iii.b)** this is associated with divergence in the quality and magnitude of functional trait responses.

## 2 Material & Methods

### 2.1 Study Species and Study Site

*Cakile maritima* is an annual herb with slightly succulent, lobed leaves and a branched shoot system that can reach ~ 1 m of lateral expansion. Its predominantly insect-pollinated flowers develop two-segmented siliques, which are dispersed by water and wind. The species inhabits the foreshores and primary dunes in marine coastal ecosystems where it is exposed to pronounced daily and seasonal variation in nutrient availability, salinity, substrate stability, soil humidity and temperature regimes (He and Silliman, 2019). *Cakile* is essential for the formation of primary dunes as well as dune succession and provides food for a broad range of pollinating and herbivorous insects (reviewed in Arbelet-Bonnin *et al*. 2019). Moreover, it is considered a promising future crop for saline agriculture aiming at the production of edible oil, biofuels and diverse bioactive compounds for pharmaceutical use, as well as on the remediation of salt and heavy metal affected soils (reviewed in Arbelet-Bonnin *et al*. 2019).

*Cakile* is native to Eurasia and expanded its distribution range to North America, Australia and New Zealand in the past centuries. The species has diversified across this wide range of latitudes with climates ranging from arid to humid leading to the sub-division in various controversial sub-species based on differences in fruit and leaf morphology (Davy et al., 2006; ITIS, 2022), physiology (e.g., Megdiche *et al*. 2008) and salt stress tolerance (reviewed in Arbelet-Bonnin *et al*. 2019). However, divergence in the ability to resist combined heat and drought stress has been largely overlooked (but see Jdey *et al*. 2014; Ellouzi *et al*. 2017 for single stress responses). In our study, we focus on the native Western-Central European distribution range of *Cakile*, which spans gradients of increasing temperatures and decreasing precipitation from northern to southern latitudes (**Tab. 1**).

**Tab. 1.**
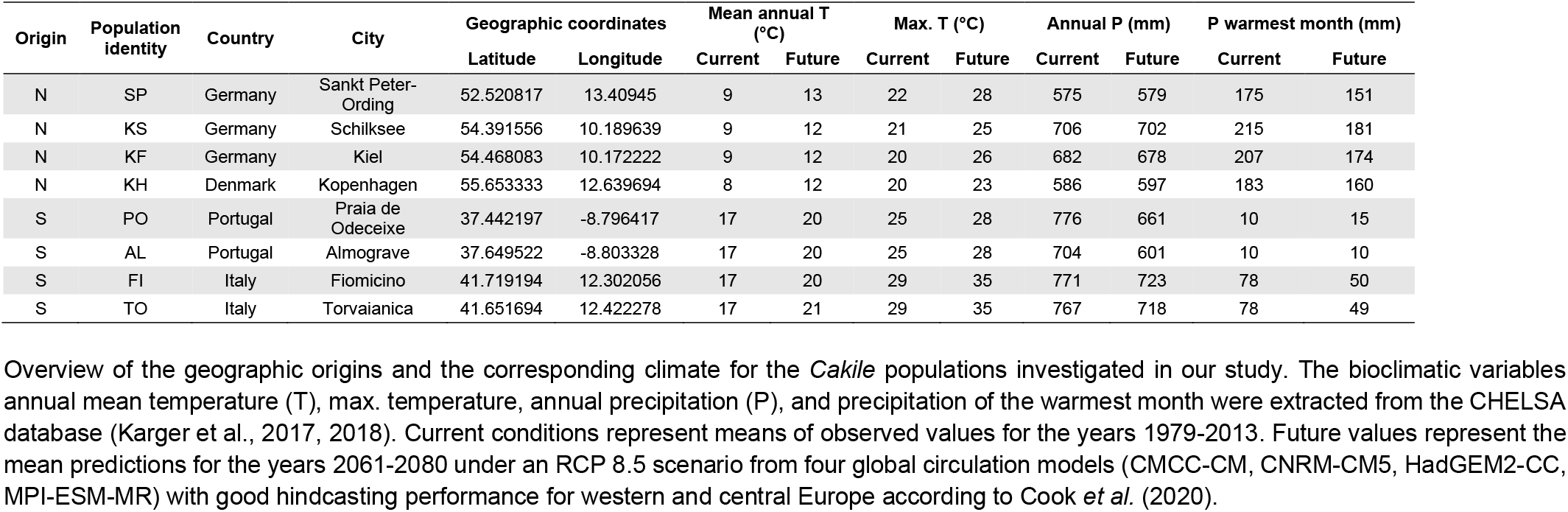
Background information on the studied populations.

| Origin | Population identity | Country | City | Geographic coordinates |  | Mean annual T (°C) |  | Max. T (°C) |  | Annual P (mm) |  | P warmest month (mm) |  |
| --- | --- | --- | --- | --- | --- | --- | --- | --- | --- | --- | --- | --- | --- |
|  |  |  |  | Latitude | Longitude | Current | Future | Current | Future | Current | Future | Current | Future |
| N | SP | Germany | Sankt Peter-Ording | 52.520817 | 13.40945 | 9 | 13 | 22 | 28 | 575 | 579 | 175 | 151 |
| N | KS | Germany | Schilksee | 54.391556 | 10.189639 | 9 | 12 | 21 | 25 | 706 | 702 | 215 | 181 |
| N | KF | Germany | Kiel | 54.468083 | 10.172222 | 9 | 12 | 20 | 26 | 682 | 678 | 207 | 174 |
| N | KH | Denmark | Kopenhagen | 55.653333 | 12.639694 | 8 | 12 | 20 | 23 | 586 | 597 | 183 | 160 |
| S | PO | Portugal | Praia de Odeceixe | 37.442197 | -8.796417 | 17 | 20 | 25 | 28 | 776 | 661 | 10 | 15 |
| S | AL | Portugal | Almograve | 37.649522 | -8.803328 | 17 | 20 | 25 | 28 | 704 | 601 | 10 | 10 |
| S | FI | Italy | Fiomicino | 41.719194 | 12.302056 | 17 | 20 | 29 | 35 | 771 | 723 | 78 | 50 |
| S | TO | Italy | Torvaianica | 41.651694 | 12.422278 | 17 | 21 | 29 | 35 | 767 | 718 | 78 | 49 |

Overview of the geographic origins and the corresponding climate for the *Cakile* populations investigated in our study. The bioclimatic variables annual mean temperature (T), max. temperature, annual precipitation (P), and precipitation of the warmest month were extracted from the CHELSA database (Karger et al., 2017, 2018). Current conditions represent means of observed values for the years 1979-2013. Future values represent the mean predictions for the years 2061-2080 under an RCP 8.5 scenario from four global circulation models (CMCC-CM, CNRM-CM5, HadGEM2-CC, MPI-ESM-MR) with good hindcasting performance for western and central Europe according to Cook *et al.* (2020).

### 2.2 Plant Propagation

Fruits were collected in accordance with their natural dispersal period between September and November 2020 in four Northern and four Southern European populations (**Tab. 1**). In each population, we collected ripe fruits from each of 10 vigorous plant individuals (i.e., maternal families), which were spatially separated between 20-50 m. The fruits were cleaned and stored in slightly damp sand at 4°C for cold stratification until January 2021. Subsequently, seeds were removed from their fruit coats, imbibed in tab-water on filter paper in petri dishes for nine days, and germinated in sand (0-2 mm grain size Min2C Kinderspielsand, Bauhaus, Mannheim, GE) at 22°C / 10°C in a 16 h light / 8 h dark cycle (MLR-352-PE equipped with FL40SS fluorescent tubes, Panasonic Health, Kadoma, JP).

Individual seedlings were pricked into 1.1 L pots (11 × 11 × 12 cm, Göttinger, Göttingen, GE) containing the above-mentioned sand and grown in climate chambers (Fitotron Typ HGC 1514 Module 7, Weiss Umwelttechnik GmbH, GE; equipped with Valoya C90 NS12 + PSU LEDs, Valoya, FIN) under standardized daily courses of temperature (max. 22, min. 10°C) and light (max. 900, min 0 μmol/m^2^s) at constant water vapour pressure deficit (VPD) of 0.85 +/0.15 kPa (**Fig. 1b** control climate conditions). During the growing phase, the plants were randomized in a two to three-week rhythm within and among climate chambers and received tab-water according to demand as well as 150 mL of a fertilizer solution (1 g/L Universol Yellow, ICL Speciality Fertilizers Germany & Austria, Nordhorn GE; 0.4 g/L sea salt, Esco European Salt Company, Hannover, GE) every two weeks.

**Fig. 1.**
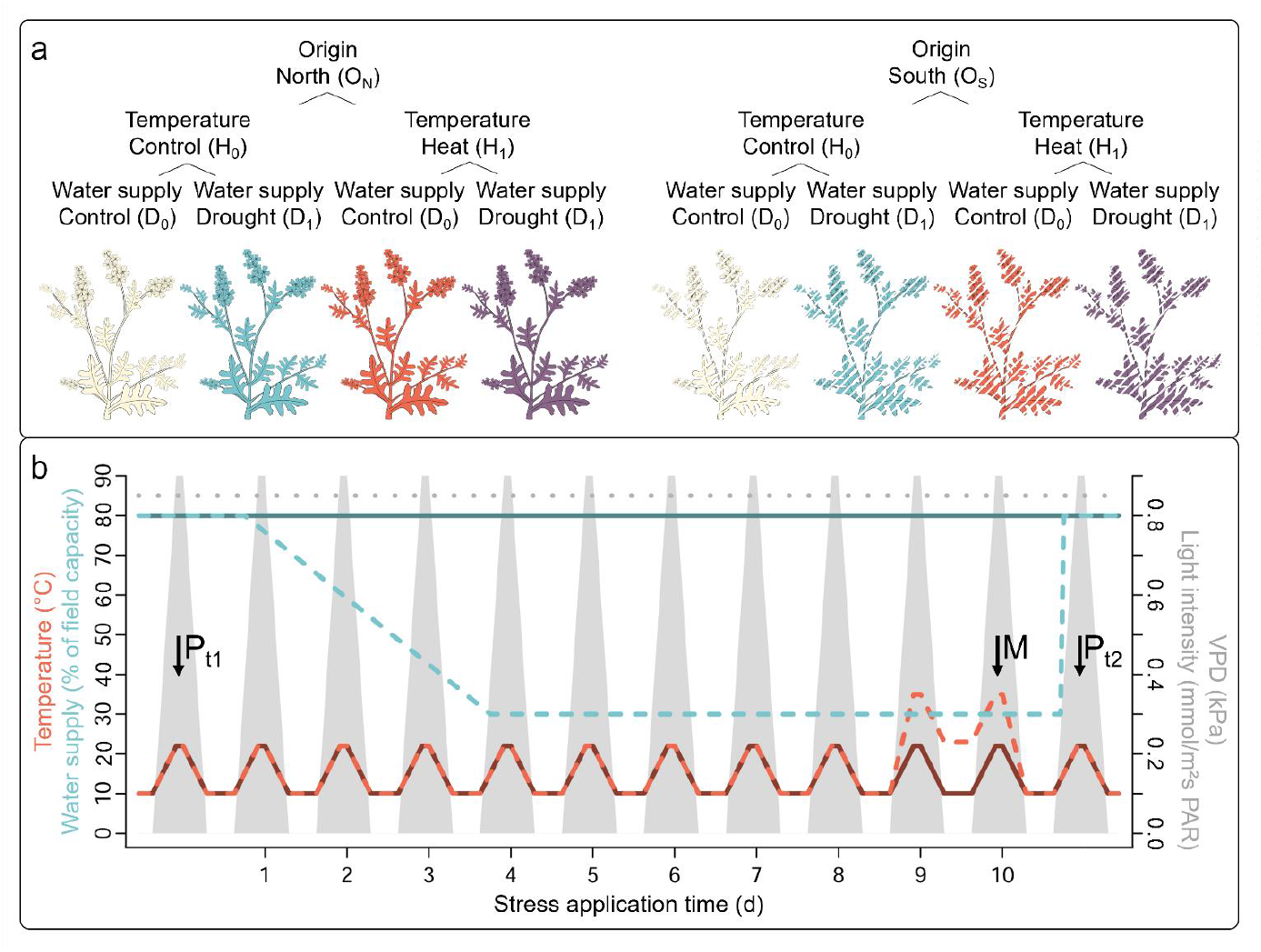
Overview of the experimental setup. **a)** Plants from four Southern and four Northern European *Cakile* populations were exposed to drought and heat stress in a fully factorial experimental design. **b)** Climate settings for the ten-day stress application. Water control plants (D_0_, solid dark blue line) were re-watered to 80% of soil field capacity every day in the morning, whereas drought stress plants (D_1_, dashed light blue line) were re-watered to only 30%. All plants assigned to the D_1_ treatment had reached 30% of field capacity at day 4. Temperature control plants (H_0_, solid dark red line) were exposed to max. 22°C and min. 10°C, whereas heat stress plants (H_1_, dashed light red line) were exposed to max. 35°C and min. 23 °C for the last two days of stress application. Plants from all treatments were grown under standardized daily courses of light (grey faces) at day lengths of 15 h with max. 0.9 mmol/m^2^s and constant VPD of 0.85 +/- 0.15 kPa (dotted grey line). The climate conditions in the plant propagation phase corresponded to control conditions for water and temperature treatment (dashed blue and red line) and the treatment-independent light and VPD regimes illustrated here. The time points for the harvest of leaf material for metabolomics (M) and the acquisition of plant phenotype data (P) are indicated by vertical arrows.

### 2.3 Experimental Setup

At the age of four months, we exposed the plants to stress. Our fully factorial experimental design comprised eight latitudinal origin (O_S_ = south, O_N_ = north) × water treatment (D_0_ = control, D_1_ = drought) × temperature treatment (H_0_ = control, H_1_ = heat) combinations, which were replicated across four *Cakile* populations with a maximum of seven maternal families per population depending on the availability of vital seedlings. This setup resulted in a total number of 165 plants for the experiment (**SI. 1**). With the treatments described in the following, we aimed at assessing differences among Southern and Northern European origins in the plants’ abilities to acclimate to drought and heat stress over short term (**Fig. 1**).

Drought stress was applied with a gravimetric procedure. Before the pots were planted with *Cakile* seedlings, we lined them with pristine sheep wool fleece (4 mm thick, Herrmann Meyer KG, GE) to prevent the loss of substrate *via* the bottom holes, filled them with an exact amount of 1.33 kg dried (24 h at 105°C) sand, and determined the mass of the entire experimental unit (*m _pot & sand at 0% FC_*). Afterwards, we watered the soil to field capacity and redetermined the experimental units’ masses (*m _pot & sand at 100% FC_*). One week prior to the application of the drought stress treatment, we watered the pots with grown plants to field capacity again to determine their mass a third time (*m _plant, pot & sand at 100% FC_*). The treatment was applied by re-watering the pots to either 80 % (control, D0) of field capacity or 30 % (drought, D1) of field capacity. The target mass of experimental units for the gravimetric water adjustment was calculated as *m _target_* = (*m _plant, pot & sand at 100% FC_* – *m _water_*) + *x* * *m _water_*; where *m _water_* is the mass of water in a given experimental unit at field capacity = *m _pot & sand at 100% FC_* – *m _pot & sand at 0% FC_*, and *x = 0.8 for D_0_, 0.3 for D_1_*. We applied the gravimetric water adjustment for ten days in the morning hours, whereby the soil water content of all plants assigned to the D_0_ treatment had fallen to 30 % of field capacity at the fourth day.

Heat stress was applied for a short period of two days at the end of the drought stress period *via* the climate chamber settings (**Fig. 1b**). Plants were exposed to either max. 22°C and min 10°C (control, H_0_) or max. 35°C and min 23°C (heat, H_1_), which is in accordance with the temperature conditions predicted by global circulation models (**Tab. 1**). The temperatures in H_0_ and H_1_ were not applied as consistent blocks but exhibited near-natural temporal dynamics with daily max. and min. temperatures (Δ = 12°C), which merged smoothly (**Fig. 1b**). At constant relative air humidity (RH), an increase in temperature automatically increases VPD and thereby stomatal transpiration (Georgii *et al*. 2017). To avoid that the heat stress treatment automatically increases the intensity of drought stress thereby making it impossible to disentangle the effects of both stressors, the air VPD in H_0_ and H_1_ treatment was kept at a constant low value of 0.85 +/- 0.15 kPa by adjusting RH (35°C ~ 85%, 22°C ~ 65%). H_0_ and H_1_ climate conditions were replicated across two chambers, within which plants were fully randomized with regard to origin, population and water treatment.

### 2.4 Data Acquisition

#### 2.4.1 Plant Growth and Morphological Functional Traits

To assess differences in plant growth responses to stress among Southern and Northern European *Cakile* populations, we counted the number of fully developed leaves and measured the cumulative shoot length, i.e., the summed length of the main shoot and all side shoots. Moreover, we counted the number of flower buds and fully opened flowers. All these traits were acquired twice directly prior to and after the stress application (**Fig. 1b**). We subtracted the values assessed prior to stress application from the values assessed after the stress application for further analyses.

All morpho-functional traits were assessed once after the stress application. We determined the number of abscised leaves (yellow, red or brown colour, necrotic). Moreover, we picked the youngest fully developed intact leaf from one randomly chosen shoot per plant to determine its fresh leaf area (Expression 11000 XL Scanner, Seiko Epson, JP; WinFOLIA™ Pro 2015, Regent Instruments, CAN) and dry mass (24 h at 101°C). Based on these proxies, we calculated the specific leaf area (SLA) as *fresh leaf area/dry leaf mass*, leaf dry matter content (LDMC) as *dry leaf mass/fresh leaf mass*, and leaf succulence as *fresh leaf mass/fresh leaf area*. In addition, we determined the above and belowground dry biomass (60 h at 101°C) for each of the plants to assess their root-shoot ratios (RSR).

#### 2.4.3 Metabolome Traits

We harvested the youngest, fully developed leaf from three randomly chosen shoots per plant individual at the last day of stress application between 1 and 2 pm, i.e., one hour after the air temperature has reached its daily maximum (**Fig. 1b**). The samples (three leaves per plant individual pooled) were immediately snap-frozen in liquid nitrogen and stored at −80°C. Hydrophilic and lipophilic leaf metabolites were extracted by a modified liquid-solid extraction in according to Matyash *et al*. (2008). About 10 mg of lyophilized and homogenized leaf powder were suspended in 500 μL ultra-pure methanol (Carl Roth, GE) and 4 mL methyl-tert-butylether (LC-grade, Carl Roth, GE). Samples were incubated for 30 min at 25 rpm in an overhead shaker before phase separation was induced by adding 500 μL ultra-pure water (Carl Roth, GE). Following 10 min of incubation, samples were centrifuged at 4000 g and 4°C for 10 min and the upper lipophilic phase was collected. The remaining hydrophilic phase was used as a template to repeat the above-described protocol for the extraction of hydrophilic metabolites. Hydrophilic and lipophilic extracts were dried at 0.1 bar and 45°C (SpeedVac, Thermo Scientific, GE) and re-suspended in 500 μl ultra-pure water and methanol (50/50, v/v, with 0.1% acetic acid, ultra-pure grade, Carl Roth, GE) or in 500 μl isopropanol and chloroform (3:1, v/v with 0.1% acetic acid, both LC-grade from Carl Roth, GE), respectively.

Hydrophilic and lipophilic leaf extracts from all experimental plants were analysed *via* Fourier Transform-Ion Cyclotron Resonance-Mass Spectrometry (FT-ICR-MS, 7 Tesla Solarix-XR, Bruker, GE). Injection was conducted by a 1260 HPLC pump and an auto-sampler (Agilent, GE) without column separation with the eluent corresponding to the extract dilution eluent. Directly injected extracts underwent electro-spray ionisation in both, the positive and the negative mode. The source temperature was set to 200°C by a dry gas flow of 4 L/min and a nebulizer pressure of 1 bar. The sweep excitation power was set to 18 %. All samples were measured within the 4 M mode, with a total mass range from 65-1500 m/z and an accumulation time of 0.5 s. We used two methods for the detection of metabolites: an ultra-small molecule method (UM) for metabolites between 65-300 m/z with 292 scans and a small molecule method (SM) with a range of 90-1200 m/z and 268 scans. Detailed technical settings for both methods are summarized in **SI. 2**.

Data processing was conducted with MetaboScape 2021b (Bruker, Bremen, GE). Sum formulas for metabolic features were calculated based on a tolerated mass error of < 2 ppm and an isotopic pattern value (including the isotope fine structure ratio) < 300. To reduce false-positive results, the seven golden rules of Kind & Fiehn (2007) were used. Features without assigned sum formulas were neglected from further analyses. We created a single merged dataset on the mass intensity (i.e., semi-quantitative approach) of all remaining metabolites for further statistical analyses. If a particular metabolite was detected in both, the positive and negative mode or the SM and UM method, we only maintained its intensity values from the mode/method resulting in the larger mean across all samples. Intensities of metabolites that were detected in both, lipophilic and hydrophilic leaf extracts were summed. All metabolites, which were not abundant in at least 80 % of the samples in at least one of the eight origin × water treatment × temperature treatment combinations, were removed from the dataset for further analyses, resulting in a total number of 1966 metabolites in the final merged dataset. Annotation of these metabolites was performed with MetaboScape 2021b based on the comparison of mass spectra and m/z ratios with the KEGG database (Kanehisa et al., 2019), HMDB (Wishart et al., 2007), Lotus database (Rutz et al., 2022) and various original research papers (e.g., Rodman 1976; Radwan *et al*. 2008; Taamalli *et al*. 2015; Omer *et al*. 2016, 2019; Arbelet-Bonnin *et al*. 2020; Placines *et al*. 2020; Shams *et al*. 2010).

The statistical analyses (see chapter **2.5**) revealed a set of significantly stress-modulated metabolites, which were further investigated *via* Ultra Performance Liquid Chromatography coupled to a Triple Quadrupole Mass Spectrometer (UPLC: ACQUITY Ultra Performance LC, TQ-MS/MS: Premier XE TripleQuad, Waters, GE) to verify annotations. Chromatographic separation of metabolites in leaf extracts was performed on a C18 gravity column (1.8 μm particles, 100 × 2.1 mm, Macherey-Nagel, GE) using ultra-pure water with 0.1% acetic acid (A) and acetonitrile with 0.1% acetic acid (B) as mobile phase at a flow rate of 250 μL/min. The gradient started at 100% A for 1 min, ran within 10 min to 100% B, was held for 1 min, flushed back to 100% A within 1 min, and was held for 3 min. The main parameters for the subsequent TQ-MS/MS are summarized in **SI. 2**. Metabolites, the concentration of which was below the detection limit of UPLC-TQ-MS/MS, were analyzed via FT-ICR-MS in the MS^2^ mode. This allows the detection of single, ultra-high-resolution fragment spectra in the ultra-trace range. The MRM as well as the MS^2^ spectra were compared with entries in HMDB (Wishart et al., 2007) and MassBank (Oberacher et al., 2020) for metabolite annotation.

### 2.5 Statistical Analyses

All statistical analyses were performed in R 4.1.2. (R Development Core Team, 2021). Plant performance traits were analysed with (generalized) linear mixed effects models ((G)LMMs. All models for responses related to plant growth and functional traits were fitted with the R-package “glmmTMB” (Brooks et al., 2017) and included origin (levels = O_N_, O_S_), water treatment (levels = D_0_, D_1_), temperature treatment (levels = H_0_, H_1_), and all possible interactions among these factors as predictors. Population identity was included as random effect. Please note that the combination of the three predictors and the random effect corresponds to the maternal family level, which is the reason why maternal family is not included as a random factor (**SI. 1**). All models were adjusted (i.e., selection of appropriate data transformations, error families, link functions, zero-inflation and dispersion formulas) and validated based on diagnostic plots and tests provided in the R-package “DHARMa” (Hartig, 2021). Sum-to-zero contrasts were set on all factors for the calculation of type III ANOVA tables based on Wald-χ^2^ tests (R-package “car”, Fox & Weisberg 2018). If origin, water and/or temperature treatment were involved in significant interactions, we calculated post-hoc contrasts among the estimated marginal means of their factor levels only within levels of other factors involved in the respective interaction (R-package “emmeans”, Lenth 2020). Variance components were extracted from all models using the R-package “insight” (Lüdecke et al., 2019) and model parameter estimates were standardized for comparative illustration using the R-package “effectsize” (Ben-Shachar et al., 2020).

We used both, multivariate and targeted statistical analyses for the metabolome dataset. In a first step, we ran random forest models with the full dataset including all 1966 metabolites (R-package “randomForest”, Liaw & Wiener 2002; R-package “party”, Strobl *et al*. 2008). Both, unsupervised (plant individuals are clustered based on their metabolome without information on origin and stress treatments) and supervised (metabolite data are used to predict the group identity O_N_D_0_H_0_, O_N_D_1_H_0_, O_N_D_0_H_1_, O_N_D_1_H_1_, O_S_D_0_H_0_, O_S_D_1_H_0_, O_S_D_0_H_1_ or O_S_D_1_H_1_ of a given plant) models were calculated. The random forest classification trees, based on 2000 individual decision trees with 44 randomly selected metabolites accepted as candidates for each split. The importance of a metabolite in predicting a given group identity in the supervised random forest models was determined as mean decrease in accuracy (MDA) of the grouping prediction upon removal of the respective metabolite from the model.

A more targeted statistical approach towards the identification of stress-modulated metabolites, is the generation of volcano plots that allow the identification of key compounds with high and significant differences in abundance between control *versus* stressed plants (Cui and Churchill, 2003). However, this approach neither allows to statistically assess interactive effects of different stressors (i.e., drought × heat interactions) or differences in stress responses among origins (i.e., origin × stress interactions), nor to account for the non-independence of individuals within populations (i.e., population random effect), differences in the error distribution structure among metabolites (gaussian, poisson, zero-inflated) and variance inhomogeneity. As such, we employed an alternative statistical approach resting on Georgii *et al*. (2017). In a first step, we calculated the fold change in the intensity of each metabolite for all origin × water treatment × temperature treatment combinations as *mean intensity in D_1_H_0_, D_0_H_1_, and D_1_H_1_ plants / mean intensity in D_0_H_0_ plants*. Metabolites with a fold change of either ≤ 0.5 (corresponds to halving of intensity under stress) or ≥ 2 (corresponds to doubling of intensity under stress) were maintained for the following statistical analyses (341 metabolites), whereas the rest was neglected. In a next step, we assessed the combined effects of origin, drought and heat with GLMMs as described above for the plant growth traits. We created R-loops for both, the model checking procedure and the effect assessment. For model checking, the 321 metabolites ran through a loop in which they were fitted as responses in models with the basic structure ~ *origin * water treatment * temperature treatment + (1/population*). Each metabolite ran with 13 alternative models differing in their response variable transformation, error families, zero-inflation formula, and dispersion formula (see **SI. 3**). We extracted the results from the nonparametric dispersion test (i.e., residuals *versus* fitted), Kolmogorov-Smirnov test for uniformity, and Levene’s test for homogeneity of variance, which are provided for targeted model optimization in the R-package “DHARMa” (Hartig, 2021), to select the best fit model. In a second loop, the best fit models for each metabolite were analysed with type III ANOVAs based on Wald-χ^2^ tests (R-package “car”, Fox & Weisberg 2018) and the calculation of post-hoc contrasts (R-package “emmeans”, Lenth 2020) as described for performance traits. The p-values from ANOVA tables were corrected for inflation of alpha errors by multiple testing (False Discovery Rate, FDR) according to Benjamini & Hochberg (1995).

## 3 Results

### 3.1 Effects of Origin, Heat and Drought on Plant Growth and Morpho-Functional Traits

*Cakile* populations from Northern and Southern European origins exhibited pronounced differences in growth habit, independently from stress treatments (**Tab. 2, Fig. 2a-e**). Plants from northern origins developed more expansive shoot systems (p = 0.016, χ^2^ = 5.818) and produced more open flowers (p = 0.003, χ^2^ = 8.581) than southern plants but had lower total biomasses (p < 0.000, χ^2^ = 15.574) as well as fewer (p = 0.006, χ^2^ = 7.654), thinner (i.e., higher SLA, p = 0.026, χ^2^ = 4.977) and less succulent (p = 0.006, χ^2^ = 7.472) leaves.

**Tab. 2.** Results of statistical plant growth analysis.

|  | **Δ Number**<br>**flower buds**<br>GLMM<br>(-, NB, -) | **Δ Number**<br>**flowers**<br>GLMM<br>(-, NB, -) | **Δ Flower**<br>**diameter**<br>LMM<br>(-, G, -) | **Δ Number**<br>**leaves**<br>GLMM<br>(-, G, -) | **Δ Shoot**<br>**length**<br>LMM<br>(-, G, -) | **Biomass**<br>LMM<br>(-, G, DF) | **SLA**<br>LMM<br>(Log, G, DF) | **LDMC**<br>LMM<br>(Srt, G, -) | **Leaf**<br>**succulence**<br>LMM<br>(Log, G, -) | **Abcised**<br>**leaves**<br>GLMM<br>(-, BB, DF) | **RSR**<br>GLMM<br>(-, BB, DF) |
| --- | --- | --- | --- | --- | --- | --- | --- | --- | --- | --- | --- |
| Intercept | 4.38 | 3.26 | -0.01 | -0.04 | 0.00 | -0.05 | 0.30 | 1.48 | 1.01 | -2.10 | -0.60 |
| Origin (O<sub>N</sub>-O<sub>S</sub>)<sub>1DF</sub> | **0.12\*\*** | **0.21\*\*** | 0.04 | **-0.22\*\*** | **0.36\*** | **-0.27\*\*\*** | **0.05\*** | 0.01 | **-0.08\*\*** | **0.24\*\*\*** | 0.00 |
| Water (D<sub>0</sub>-D<sub>1</sub>)<sub>1DF</sub> | **0.09\*\*\*** | -0.02 | 0.14 | 0.14 | **0.17\*\*** | **0.22\*\*** | **0.03\*** | -0.03 | -0.03 | -0.09 | -0.07 |
| Temperature (H<sub>0</sub>-H<sub>1</sub>)<sub>1DF</sub> | 0.02 | -0.03 | **0.31\*\*\*** | -0.14 | -0.03 | -0.12 | 0.01 | 0.02 | -0.02 | 0.01 | -0.03 |
| Origin × water<sub>1DF</sub> | **0.05\*** | 0.02 | -0.06 | 0.02 | 0.01 | 0.10 | -0.01 | -0.01 | 0.01 | **0.12\*** | **-0.10\*\*** |
| Origin × temperature<sub>1DF</sub> | 0.00 | -0.02 | 0.05 | -0.14 | -0.08 | -0.01 | -0.01 | 0.00 | 0.01 | 0.01 | 0.05 |
| Water × temperature<sub>1DF</sub> | 0.00 | 0.04 | 0.00 | -0.05 | 0.00 | 0.00 | 0.00 | -0.01 | 0.02 | 0.03 | 0.01 |
| Origin × water × temperature<sub>1DF</sub> | 0.01 | -0.01 | 0.14 | 0.05 | 0.05 | -0.08 | 0.00 | -0.01 | 0.01 | -0.06 | 0.01 |
| Variance fixed | 0.20 | 0.14 | 0.14 | 0.10 | 0.17 | 0.15 | 0.10 | 0.02 | 0.18 | 0.04 | 0.02 |
| Variance random | 0.05 | 0.07 | 0.00 | 0.00 | 0.14 | 0.00 | 0.06 | 0.00 | 0.11 | 0.00 | 0.00 |
| Variance residuals | 0.76 | 0.80 | 0.86 | 0.90 | 0.69 | 0.85 | 0.84 | 0.98 | 0.72 | 0.96 | 0.98 |

Results from the (generalized) linear mixed effects models ((G)LMM) assessing the individual and combined effects of origin, water treatment and temperature treatment on the performance of *Cakile*. The table provides standardized parameter estimates for predictors with their significance levels assessed based on Wald- $\chi^2$ -tests (\* $p < 0.05$ , \*\* $p < 0.01$ , \*\*\* $p < 0.001$ ) as well as variance components for each of the models. Significant effects are printed in bold. Predictors are listed with their factor levels (first level mentioned corresponds to intercept) and numerator degrees of freedom. Response variables are listed with the applied transformations (- = none, log = natural logarithm, srt = square root), the fitted error families (BB = Betabinomial, G = Gaussian, NB = Negative Binomial, i.e., over-dispersed Poisson with quadratic parametrization) and information on whether or not a dispersion formula was included to account for variance inhomogeneity (- = no dispersion formula included, DF = dispersion formula ~ origin × water treatment × temperature treatment).

**Tab. 3.**
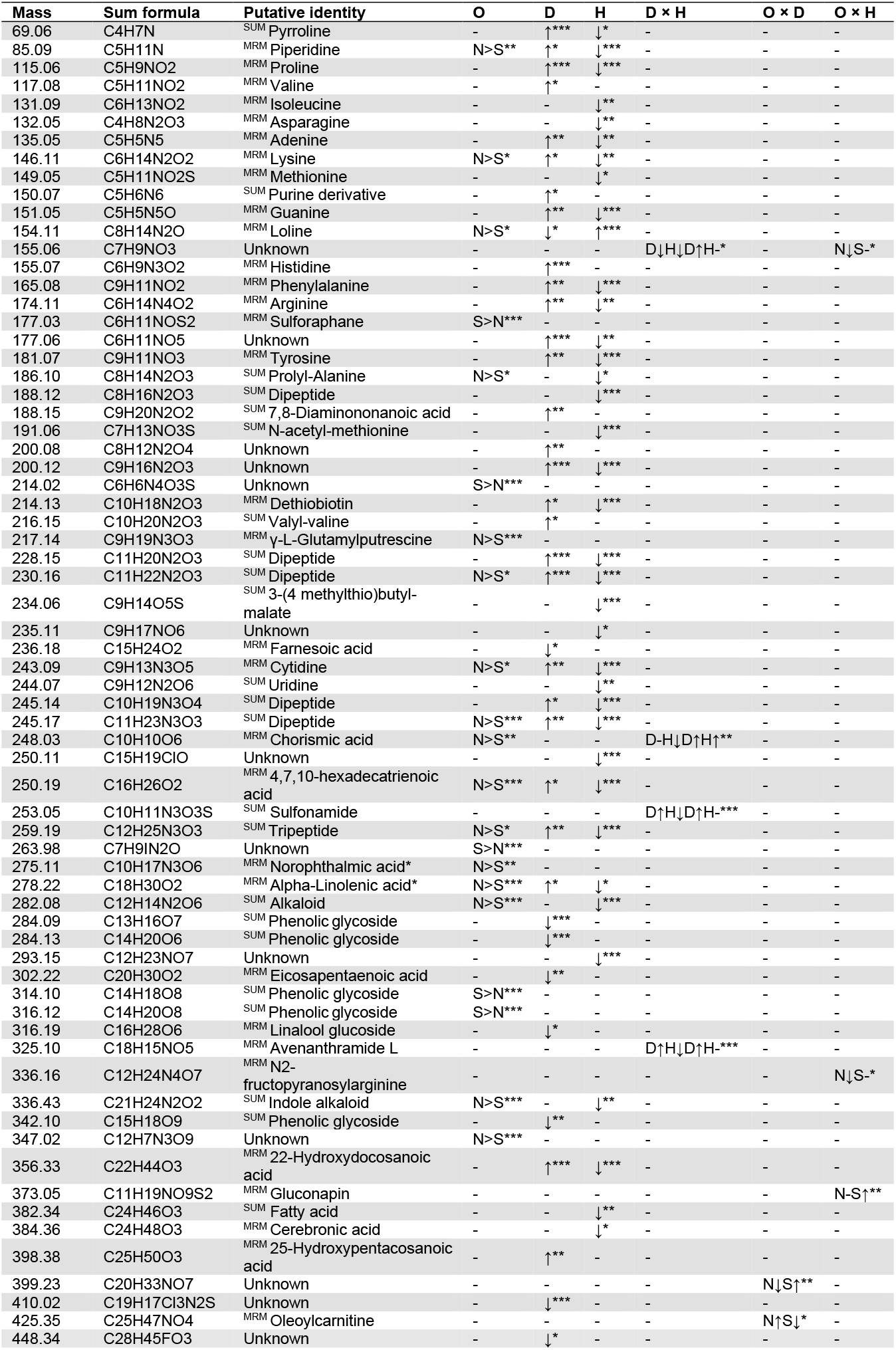
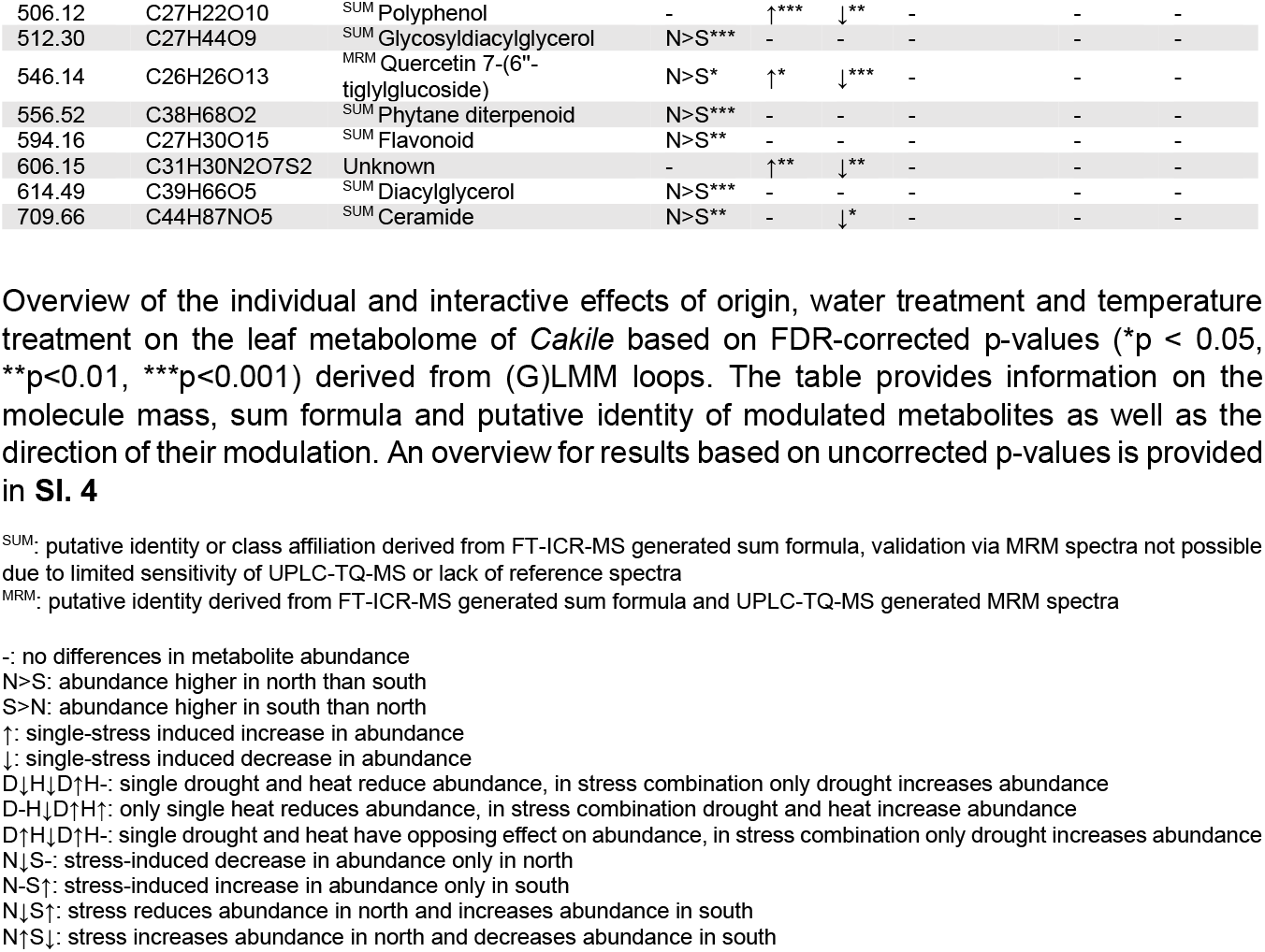
Results of targeted statistical metabolome analysis.

**Fig. 2.**
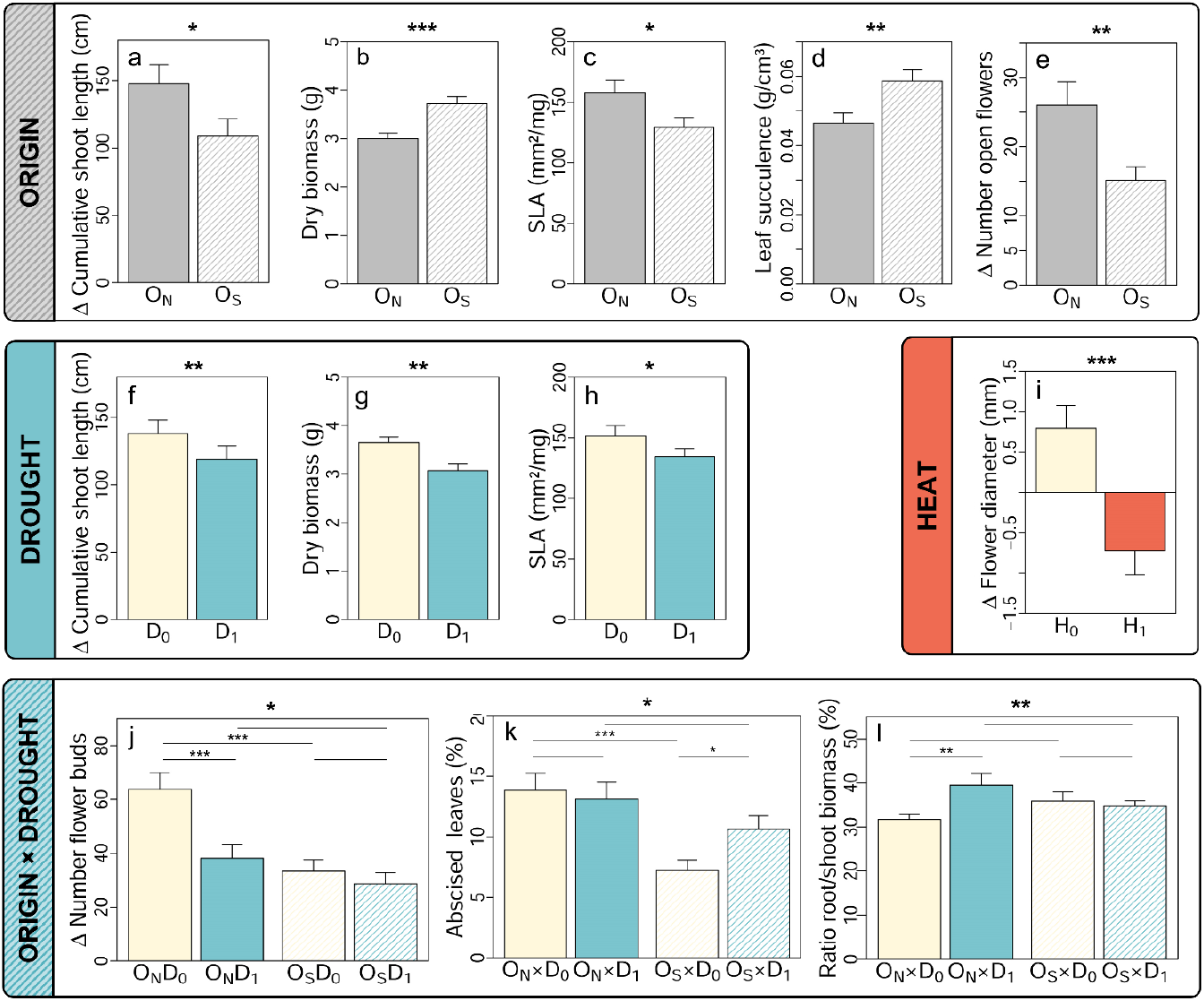
Results of plant growth analyses. Individual and combined effects of origin (North [O_N_] = filled, South [O_S_] = hatched), water treatment (control [D_0_] = cream, drought [D_1_] = blue), and temperature treatment (control [H_0_] = cream, heat [H_1_] = red) on the performance of *Cakile*. Bars represent estimated marginal means with standard errors predicted by (G)LMMs. Significance levels of main and interaction effects as assessed by Wald-χ^2^ test are denoted at the bottom of each plot (*p < 0.05, **p<0.01, ***p<0.001). Horizontal lines with asterisks within plots highlight results from post-hoc comparisons. Each origin × water treatment × temperature treatment combination was represented with 17-24 replicates. Please note, that non-destructive growth traits were acquired twice, after and before the stress application and that we refer to the difference in their values (Δ) among time points here.

Moreover, stress affected the performance of *Cakile* partially independently from population origin (**Tab. 2**, **Fig. 2f-i**). Drought stress reduced shoot length (p < 0.007, χ^2^ = 7.326), total biomass (p = 0.002, χ^2^ = 9.801) and SLA (p = 0.026, χ^2^ = 4.978), while heat stress decreased flower diameter (p < 0.000, χ^2^ = 15.172) in plants from both, northern and southern origins. Interactive effects of drought and heat stress on plant performance and functional traits were not detected (**Tab. 2**).

While the effects of heat stress were fully independent of plant origin, drought stress and origin exhibited significant interactive effects on the production of flower buds (p = 0.021, χ^2^ = 5.346), leaf abscission (p = 0.040, χ^2^ = 4.215), and root-shoot ratios (p < 0.010, χ^2^ = 6.661) (**Tab. 2**, **Fig. 2j-l**). The number of flower buds was significantly reduced by drought stress only in plants from northern origins (O_N_: t-ratio_D0-D1_ = 4.120, p_post_ < 0.000, O_S_: t-ratio_D0-D1_ = 1.001, p_post_ = 0.317) and as such northern plants only produced significantly more flower buds than southern plants when not exposed to drought stress (D_0_: t-ratio_ON-OS_ = 4.118, p_post_ < 0.000, D_1_: t-ratio_ON-OS_ = 1.453, p_post_ = 0.148). The proportion of abscised leaves increased significantly under drought stress only in Southern European populations (O_N_: t-ratio_D0-D1_ = 0.374, p_post_ = 0.709, O_S_: t-ratio_D0-D1_ = −2.514, p_post_ = 0.013). At the same time, senescent leaves were more abundant in northern plants, with a significant effect size only for drought control conditions (D_0_: t-ratio_ON-OS_ = 4.191, p_post_ < 0.000, D_1_: t-ratio_ON-OS_ = 1.411, p_post_ = 0.160). Finally, an increase in root to shoot ratio under drought stress was only detected in plants from northern origins (D_0_: t-ratio_ON-OS_ = −2.991, p_post_ = 0.003, D_1_: t-ratio_ON-OS_ = 0.501, p_post_ = 0.617).

### 3.3 Effects of Origin, Drought and Heat on the Plant Metabolome

#### 3.3.2 Multivariate Statistical Analyses

The unsupervised random forest model for the full dataset revealed no apparent clustering of plant individuals among plant origins and stress treatments based on the composition of the metabolome (**Fig. 3a**). The supervised model had only 39% accuracy in predicting a given origin × water treatment × temperature treatment combination from leaf metabolites (out of bag error = 61%), whereby false predictions were mainly made among stress treatments within origins but less among origins (**Fig. 3b**, coloured bars). This finding supports that plant origin alone explains more variation in the overall metabolome as compared to stress treatments. Accordingly, we calculated a second supervised random forest model to predict origin only from the leaf metabolome, which indeed had 93% accuracy (**Fig. 3b**, grey bars). The 20 most decisive metabolites for the correct prediction of plant origin included 10 organo-sulphur compounds, of which 6 were glucosinolates or isothiocyanates (**Fig. 3c**). Some of the latter were more abundant in northern plants (e.g., sinigrin), whereas others were more abundant in southern plants (e.g., sulforaphane, iberin, and hesperin). In summary, the results from supervised random forest models revealed that plants from northern and southern origins represent different glucosinolate chemotypes.

**Fig. 3.**
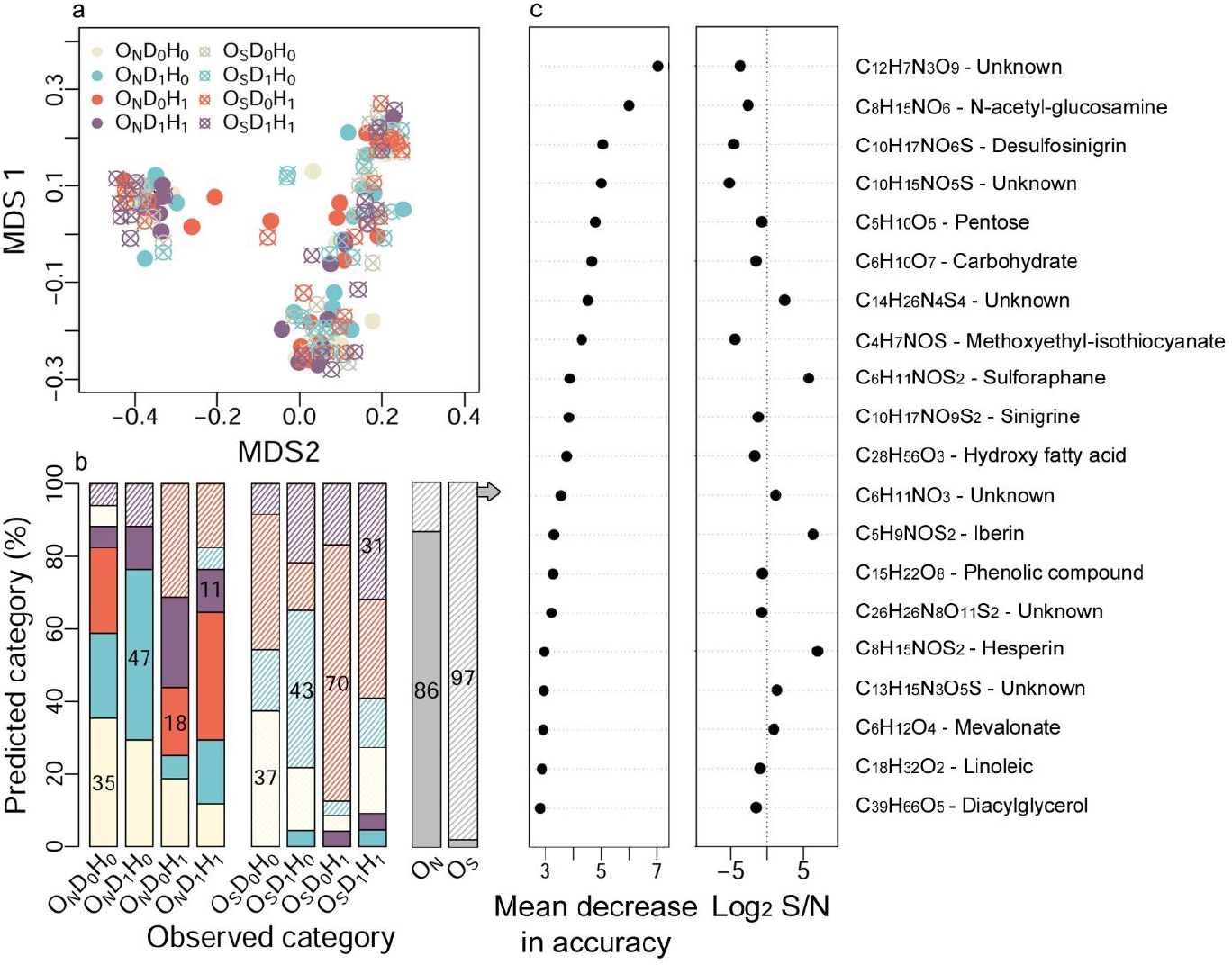
Results of multivariate statistical metabolome analyses. Combined effects of plant origin (North [O_N_] = filled, South [O_S_] = hatched), water treatment (control [D_0_] = cream, drought [D_1_] = blue), and temperature treatment (control [H_0_] = cream, heat [H_1_] = red) on the composition of 1966 leaf metabolites in *Cakile* as assessed via random forest models. **(a)** Multi-dimensional scaling plot derived from the unsupervised model. **(b)** Stacked barplot illustrating the confusion matrix from two supervised models. Colored bars base on a model predicting the origin × water treatment × temperature treatment category of plants from their leaf metabolome, while gray bars base on a model predicting only origin. Plots quantify the fraction of plants that was predictively assigned to a given category. Percentage values for the fraction of correct predictions in each observed category are denoted within the correspondingly colored block. **(c left)** Importance plot indicating 20 metabolites, the removal of which from the supervised model for plant origin causes the highest mean decrease in accuracy. **(c right)** Log_2_-fold difference in the abundance of the regarding metabolites among southern and northern origins. Negative values indicate higher metabolite abundance in northern plants, positive values higher abundance in southern plants.

#### 3.3.3 Targeted Statistical Analyses

To address differences in metabolic stress-responses among origins more specifically, we performed targeted statistical analyses with (G)LMMs on a reduced data set including exclusively metabolites with log_2_-fold changes >1 and >-1 across stress treatments. Around 92% of the observed effects origin, drought and heat were statistically additive (**Tab. 4**, **Fig. 4**). We identified 27 metabolites, which differed significantly between plants from northern and southern origins, independently from both stress treatments. Five of these metabolites were more abundant in southern than northern populations, while the remaining 22 had higher intensities in northern populations (**Tab. 4**). Southern plants had higher amounts of the isothiocyanate sulforaphane (**Fig. 4a**) as well as another not further identified sulphur-containing metabolite (C_6_H_6_N_4_O_3_S) and exhibited a higher abundance of two phenolic glycosides (C_14_H_18_O_8_, C_14_H_20_O_8_). Northern plants produced higher amounts of the alkaloids piperidine and loline (**Fig. 4b,c**), which were verified via molecule masses, sum formulas and MRM spectra; and two not further identified alkaloid-like compounds (C_12_H_14_N_2_O_6_, C_21_H_24_N_2_O_2_). Moreover, northern plants exhibited higher abundances of quercetin and another not further identified flavonoid (C_27_H_30_O_15_), several dipeptides, and lipid compounds.

**Fig. 4.**
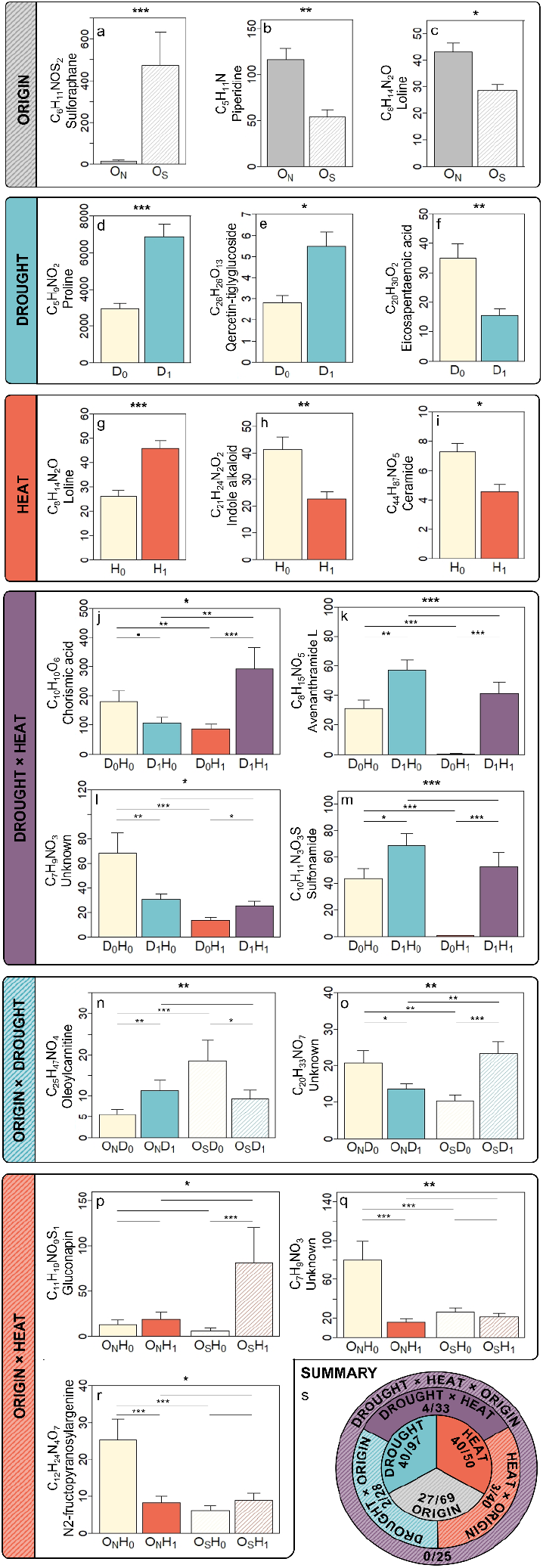
Results of targeted statistical metabolome analyses. Individual and combined effects of plant origin (North [O_N_] = filled, South [O_S_] = hatched), water treatment (control [D_0_] = cream, drought [D_1_] = blue), and temperature treatment (control [H_0_] = cream, heat [H_1_] = red) on the composition of leaf metabolites in *Cakile*. Barplots **(a-o)** show marginal estimated means for metabolite intensities (× e^06^) with standard errors predicted by (G)LMMs. Significance levels of main and interaction effects as assessed by Wald-χ2 test are denoted at the bottom of each plot (based on FDR-corrected p-values), while horizontal lines with asterisks within plots highlight results from post-hoc comparisons (• p> 0.07, *p < 0.05, **p<0.01, ***p<0.001). Each origin × water treatment × temperature treatment combination was represented with 17-24 replicates. **(p)** Provides a summary for the number of metabolites, which were significantly modulated by a given main or interaction effect as noted within the corresponding surface area based on FDR-corrected p-values / uncorrected p-values. A complete overview of results based on FDR-corrected / uncorrected p-values is provided in **Tab. 4 / SI. 4**.

Most drought and heat stress effects on the leaf metabolome of *Cakile* were independent from each other as well as from plant origin (**Tab. 4**). Drought significantly increased the intensity of 31 compounds including numerous amino acids such as proline (**Fig. 4 d**), various dipeptides, purines and pyrimidines, unsaturated long chain fatty acids, the jasmonic acid precursors 4,7,10-hexadecatrienoic acid and α-linolenic acid, and flavonoids such as quercetin (**Fig. 4 e**). The amounts of nine metabolites including loline, farnesoic acid, linalool glycoside and eicosapentaenoic acid (**Fig. 4 f**), as well as three phenolic glycosides (C_13_H_16_O_7_, C_14_H_20_O_6_, C_15_H_18_O_9_) were decreased in response to drought. Heat stress induced a significant increase in the abundance of a single compound - the alkaloid loline (**Fig. 4.g**) - but reduced the abundance of 39 metabolites including other alkaloids and alkaloid-like compounds (e.g., piperidine, C_12_H_14_N_2_O_6_, C_21_H_24_N_2_O_2_), amino acids, dipeptides, and lipids such as a ceramide (**Fig. 4h**) and α-linolenic acid. Most of the mentioned metabolites were additively modulated by heat and drought, whereby the direction of modulation was always opposed. Around 77% of the metabolites that increased in abundance under drought were simultaneously reduced by heat (**Tab. 4**). However, we also detected five metabolites, which were interactively modulated by drought and heat. Chorismic acid was reduced by individually applied drought and heat but significantly increased by both stressors in the presence of the other one (**Fig. 4j**). The alkaloid avenanthramide (**Fig. 4h**) and a not further identified sulfonamide (C_10_H_11_N_3_O_3_S) (**Fig. 4m**) were strongly decreased by heat only if plants were not simultaneously affected by drought and increased under drought in both temperature treatments. Another unidentified compound (C_7_H_9_NO_3_) was reduced by individual drought and individual heat but increased under drought in the H1 treatment (**Fig. 4l**).

Most importantly, we observed interactive effects of origin with drought and/or heat (**Fig. 4n-r**). Oleoylcarnitine had higher constitutive abundance in southern than northern plants and was drought-reduced in the former but drought-decreased in the latter, which deflated origin-specific differences in the drought stress treatment (**Fig. 4n**). In contrast, the abundance of the unknown metabolite C_20_H_33_NO_7_ was reduced by drought stress in northern plants but increased in plants from Southern European origins (**Fig. 4o**). Compared to southern origins, northern plants had higher amounts of this compounds in the absence of drought but lower amounts under drought stress. Moreover, the abundance of the glucosinolate gluconapin increased significantly in response to heat stress only in plants from southern origins (**Fig. 4n**). N2-Fructopyranosylargenine (**Fig. 4p**) and the unknown metabolite C_7_H_9_NO_3_ (**Fig. 4o**) had higher constitutive abundance in northern than southern plants but were reduced under heat in northern plants to the abundance level of southern plants. Finally, it should be noted that numerous of such interactive effects existed when neglecting FDR-correction. An overview for the results of metabolome analyses based on uncorrected p-values is provided in **SI. 4 and Fig. 4s**. Here, we observed 63% additive and 37% interactive effects of origin, water treatment and temperature treatment and it was striking that drought-induced increases of primary metabolites were largely more pronounced in southern than northern plants (e.g., proline, valine, methionine, arginine), whereas heat-induced reductions were largely stronger in northern origins (e.g., valine, leucine, lysine, methionine, arginine).

## 4 Discussion

Our study provides broad evidence for latitudinal divergence in traits associated with plant stress resistance among natural populations of *Cakile*. In the following, we discuss that **i)** plants from Northern and Southern Europe exhibit pronounced constitutive differences in growth as well as morphological and biochemical functional traits, which likely convey adaptation to local abiotic and biotic habitat conditions; **ii)** individual drought and heat trigger numerous functional trait responses in *Cakile*, while unique responses to the stress combination were only observed for biochemical traits, which highlights their relevance in plant multi-stress resistance; and **iii)** growth and functional trait responses to stress differed among plants from southern and northern origins, supporting that the former employ more efficient mechanisms to cope with heat and drought.

### 4.1 Latitudinal Divergence in Plant Growth Phenotypes and Defence Chemotypes

As expected, plants from northern and southern origins exhibited a broad set of constitutively expressed differences in growth and functional traits that likely arose from divergent selection in their local habitats. Northern populations produced less overall biomass and fewer leaves but flowered earlier (flowering plants at t1-two weeks: 85% north, 60% south) and invested more in reproductive structures such as flowers and flowering stalks that made up a huge fraction of the shoot system (**Fig. 2**). Latitudinal clines in the trade-off relationship between early flowering and investment in vegetative growth are reported for many plant species and assumed to arise from adaptation to shorter durations of the growing season towards the north (De Frenne et al., 2013; Wolkovich et al., 2014). Moreover, southern plants exhibited smaller SLA i.e., lower leaf transpiration rates and higher leaf succulence i.e., water storage (**Tab. 2**), which mediate adaptation to more frequent and intense drought events (Kooyers et al., 2015). High leaf succulence simultaneously enhances the dilution of Na^+^ and Cl^-^ ions in the cell, thereby potentially mediating adaptation to higher ocean salinities in southern populations (southern origin: Atlantic and Mediterranean seas with 3.5-3.7%, northern origins: Baltic sea and river estuaries at North sea with 0.8-2%) (de Vos et al., 2010; Zweng et al., 2019).

The plant metabolome also exhibited constitutive latitudinal divergence. The unsupervised random forest model for the full dataset revealed no apparent clustering of origins (**Fig. 3a**), which was expected, given that most leaf metabolites fulfil basic functional tasks and should thus be highly conserved within species. Nevertheless, the supervised model had high accuracy in predicting plant origin from several fatty acids, glucosinolates and isothiocyanates with southern plants having remarkably higher abundances of hesperin and sulforaphan (**Fig. 3b,c**). The targeted statistical analyses (**Tab. 2**) confirmed higher amounts of certain isothiocyanates in southern populations and additionally revealed that northern plants contain more piperidine and indole alkaloids (**Tab. 2**). In Brassicaceae species, glucosinolates, and alkaloids constitute an important part of the defence against herbivores and pathogens (Brock et al., 2006; Nafisi et al., 2006; Liu et al., 2014). Population divergence in the composition of glucosinolates was reported for other crucifers across latitudinal, longitudinal and elevational gradients (Züst et al., 2012; Buckley et al., 2019). whereas information on intra-specific differentiation in alkaloid contents in the family is scarce (but see Zhang *et al*. 2014), given that individual compounds are largely species-specific (Nafisi et al., 2006). The extensive differentiation of glucosinolate chemotypes in *Arabidopsis* is known to be mediated by polymorphisms at only few genetic loci, the allele frequencies of which can shift quickly under selection by particular herbivore species in a given geographic region (Züst et al., 2012). Likewise, De-la-Cruz *et al*. (2020) demonstrated that the local herbivore community is a primary selective agent for the divergent evolution of alkaloid chemotypes in the Solanaceae species *Datura stramonium*. Both studies stressed that it is not the total amount of glucosinolates or alkaloids, but rather their composition as characterized by particularly high abundances of a few key compounds that maximizes plant fitness in a given environment. As such, latitudinal divergence in the composition of glucosinolates and alkaloids in *Cakile* is likely driven by the local community structure of pathogens and herbivores, which remains to be tested in future studies.

Finally, we observed a significantly higher abundance of loline in northern *Cakile* plants (**Fig. 4c**, **Tab. 4**, **SI. 5**). To our knowledge, this is the first observation of loline alkaloids in Brassicaceae. These compounds are known to be abundant in Poaceae species, where they are produced by symbiotic endophytic fungi of the genus *Epichloë* to defend their host plants against herbivores and pathogens (Bastias et al., 2017). Our study provides a first indication on the occurrence of such symbiosis in Brassicaceae species and additionally supports that this interspecific interaction may exhibit geographic variation, the exploration of which opens interesting future avenues for research.

### 4.2 Plant Abiotic Stress Responses Across Organizational Levels

We expected drought and heat to reduce plant growth in an interactive manner. Indeed, flower size decreased under heat while drought reduced plant dry biomass, the expansion of the shoot system and -partially-the production of flower buds (**Fig. 2**). However, these effects were purely additive, which is likely related to the short heat stress application time and the constant VPD across temperature treatments in our experiment. Varying relative air humidities across heat treatments to achieve constant VPDs is necessary to disentangle the physiological effects of increased temperatures from water deficit (at constant air humidity, VPD and thus leaf transpiration rates and water deficit increase with temperature). However, in the field where drought and heat typically reduce growth in an interactive manner (e.g., Hoeppner & Dukes 2012), VPDs increase with temperature. As such, our findings for additive drought and heat effects on plant growth at constant VPD support the assumption, that interactive effects in the field may arise from increased drought stress intensity in the combined stress treatment (Bita & Gerats 2013; Georgii *et al*. 2017).

The reduced growth of *Cakile* under stress may either reflect a direct detrimental effect, or alternatively result from a re-allocation of resources to mechanisms that recover homeostasis and maintain fitness under stress. On the morphological level, such responses were exclusively observed under drought and consisted e.g., of an SLA reduction to lower transpiration rates. The metabolome changed comprehensively in response to both, drought and heat (**Tab. 4**). Drought increased the abundance of numerous single amino acids and their associated dipeptides including proline, which was also observed under physiologically similar salinity stress in *Cakile* (Arbelet-Bonnin *et al*. 2020). The accumulation of amino acids under osmotic stress maintains homeostasis by increasing leaf water potential, stabilizing proteins or membranes, and scavenging ROS (Hayat et al., 2012), whereby the high abundance of dipeptides indicates their active transport across plant organs (Pu et al., 2018). Moreover, drought stressed plants had higher amounts of the flavonoid quercetin and a polyphenol (C_27_H_22_O_10_) with high antioxidative potential (Anand David et al., 2016) but reduced amounts of several small phenolic glycosides, which may provide precursors for the synthesis of the former. We also observed drought-induced changes in lipid profiles with a significant decrease of unsaturated fatty acids (farnesoic acid, eicosapentaenoic acid) and an increase of unsaturated fatty acids (hydroxydocosanoic acid, hydroxypentacosanoic acid). The former changes may relate to lipid remodelling in the membranes of water-deprived cells, which maintains their stability and integrity under drought (Kumar et al., 2021), while the latter changes may associate with the thickening of leaf cuticular layers for reduced transpiration (Fernández et al., 2016). Several drought-increased metabolites such as 4,7,10 hexadecatrienoic acid or α-linolenic acid are precursors of the phytohormone jasmonic acid and may thus be related to drought stress signalling and the induction of the above-mentioned metabolic changes (Wang et al., 2020).

Notably, the only metabolite that increased in response to heat was loline, an alkaloid that is more likely produced by an endophytic fungal symbiont than by the plant itself (see chapter **4.1**). Apart from that, heat reduced the abundance of numerous amino acids and dipeptides in *Cakile*. This may either be associated with the increased synthesis of chaperone proteins (Wang et al., 2004) or a general disruption of homeostasis. The latter is further supported by the heat-induced reduction of numerous other primary and specialized metabolites. Particularly, the reduction two jasmonic acid precursors (4,7,10 hexadecatrienoic acid or α-linolenic acid) is line with previous studies reporting detrimental heat effects on plant immunity and defences (Desaint et al., 2021). At the same time, unsaturated fatty acids that increased under drought, likely in association with cuticular thickening for reduced transpiration, were reduced under heat, likely to increase transpiration. This finding underlines that individual drought and heat may foster opposing plant responses and thus require unique responses when occurring simultaneously.

Indeed, we observed interactive effects of drought and heat on the abundance of several specialized metabolites. Chorismic acid, a metabolite generated in the final step of the shikimate pathway, was moderately reduced by individual drought and heat but strongly enhanced by either stressor the presence of the other one (**Fig. 4j**), which suggest a growing need for the synthesis of aromatic amino acids, electron transport carriers, plant hormones, defence compounds, antioxidants or signalling molecules in plants exposed to multiple stressors (Lynch, 2022). Moreover, the phenolic alkaloid avenanthramide and a not further identified sulfonamide, both of which play roles in plant defence against pathogens (Okazaki et al., 2004; Pingaew et al., 2021), were strongly reduced by individually applied heat but increased under individually applied drought, whereas their abundance in the combined stress treatment was similar to control plants (**Fig. 4k,m**). An accumulation of alkaloids under drought was observed in several plant species and supports a function of these compounds in abiotic stress resistance, which is to date only poorly understood but might root in signalling and antioxidative functions (Jan et al., 2021; Yeshi et al., 2022). The heat-induced reduction of these compounds is in line with numerous studies reporting a downregulation of defences and increased disease or herbivore susceptibility in crops at high temperatures (Desaint et al., 2021; Schwarczinger et al., 2021).

In summary, the additive and interactive stress effects on the growth, morphology and metabolome of *Cakile* suggest that the species can effectively acclimate to drought stress, whereas short-term heat events seem to cause a general disruption of homeostasis.

### 4.3 Latitudinal Divergence in Morpho-biochemical Plant Responses to Drought and Heat

In accordance with our expectations, Northern and Southern European *Cakile* differed considerably in their functional trait responses to stress on the morphological (**Fig. 2j-l**) and biochemical (**Fig. 4h-r**) level. Northern plants responded to drought with an expansion of the root system to improve water uptake and simultaneously reduced investment in flower production. Hence, they employed plastic responses mediating drought stress avoidance (i.e., preventing water deficit at the cellular level), which came at the cost of reproduction. Alternatively, reduced flower production in northern plants may be interpreted as a direct negative effect of stress. In contrast, southern plants remained their investment in both, reproductive and root structures while initiating leaf abscission to recycle resources and reduce transpiration rates (Munné-Bosch and Alegre, 2004). Hence, they employed only minimal plastic growth responses to drought stress, while maintaining their reproductive function. This suggests that southern plants either have sufficient constitutive drought stress avoidance strategies, which could be manifested in lower SLAs and higher leaf succulence (see **4.1**), or that they tolerate stress, i.e., are able to maintain cellular homeostasis even under water deficit. Such tolerance strategies may manifest on the level of the metabolome.

Indeed, numerous drought-induced metabolite increases, which are known to maintain homeostasis under drought (e.g., proline) were significantly stronger in southern than northern plants when not correcting for multiple testing (**SI. 4**). Similarly, the heat-induced reductions of numerous amino acids and dipeptides were significantly more pronounced in northern than southern plants when not accounting for FDR, which suggest a more severe disruption of homeostasis in the former. While FDR-correction is necessary to statistically account for inflation of α-errors, it should not unconditionally lead to the neglection of such effects, since even moderate changes in the abundance of metabolites can have large-scale functional consequences. Most notably, several origin × stress interactions remained also after FDR-correction and further supported more efficient stress responses and less stress damage in southern plants. We observed higher constitutive amounts of acylcarnitine in southern plants with an opposite drought-stress modulation that equalized the metabolites’ abundance among origins (**Fig. 4l,m**). Oleoylcarnitine has multiple functions in plant metabolism ranging from intracellular fatty acid trafficking and energy metabolism to protection against osmotic and antioxidative stress and stress signalling (Jacques et al., 2018). As such, the higher constitutive oleoylcarnitine amounts detected in southern *Cakile* may have provided them with the opportunity to quickly initiate the latter pathways upon stress exposure. Moreover, the abundance of the isothiocyanate gluconapin increased under heat, only in southern origins (**Fig. 4p**). Elevated temperatures have been shown to enhance glucosinolate biosynthesis in several crucifers, whereby studies on *Arabidopsis* support that glucosinolates and their hydrolysis products induce the expression of heat shock proteins (HSPs) and thus substantially contribute to plant heat tolerance (Martínez-Ballesta *et al*. 2013). Finally, northern plants had higher constitutive amounts of N2-fructopyranosylarginine, which were reduced under heat to the abundance level exhibited by southern plants across both temperature treatments (**Fig. 4o,p**). Just like loline, N2-fructopyranosylargenine is known to be produced by symbiotic fungal endophytes. The genus *Alternaria* has been shown to excrete substantial amounts of this compound to promote the growth of its host plants (Wang et al., 2022). Given that *Cakile* is well known to associate with fungi of the genus *Alternaria* (Chalbi et al., 2021), our findings are indicative of a higher abundance and more severe heat-stress induced disruption of such beneficial interactions in plants from Northern European origins.

### 4.4 Conclusions and Outlook

Our study supports that local habitat conditions have fostered divergent selection on the constitutive and stress-induced expression of numerous morphological and biochemical functional traits in *Cakile* with Southern European plants exhibiting overall higher abiotic stress resistance. Given this variation, the species’ short life cycle and high rates of genetic exchange among populations (Gandour et al., 2008), *Cakile* may be capable to adapt to rapid climate change. Next to origin-specific differences in changes of root and leaf architecture, osmolytes, and antioxidants, which are well known to mediate abiotic stress resistance, we observed differences in the drought and heat-modulation of defence compounds among southern and northern plants. Our study provides indication that some of these defence compounds may not be produced by the plant itself, but by fungal endophytes. This highlights *Cakile* as a perfect model system for studying the effects of abiotic stress on biotic interactions with symbiotic and pathogenic microbiota in the context of local adaptation, which requires further controlled experiments as well as long-term reciprocal transplant experiments under near-natural field conditions.

## 0.11 Abbreviations

D_0_: Water treatment level control
D_1_: Water treatment level drought
FDR: False discovery rate
FT-ICR-MS: Fourier Transform-Ion Cyclotron Resonance-Mass Spectrometry
(G)LMM: (Generalized) linear mixed effects model
H_0_: Temperature treatment level control
H_1_: Temperature treatment level heat
LDMC: Leaf dry matter content
O_N_: Plant origin level north
O_S_: Plant origin level south
RH: Relative air humidity
RSR: Root shoot ratio
SLA: Specific leaf area
UPLC-TQ-MS/MS: Ultra Performance Liquid Chromatography-Triple Quadrupole-Mass Spectrometry
VPD: Vapour pressure deficit

## 5 Acknowledgements

We warmly thank Ute Jandt, Rossella Bellini, Ginevra Bellini, Oscar Alberto Rojas Castillo and Miguel Ángel González Pérez for collecting seeds of *Cakile*. We are grateful for the technical assistance provided by Verena Zajonc, Imke Meyer, Anne Noltenius, Marcus Schütz, Mathias Bahns, Meike Pfeiler, Julia-Jensen-Kroll and Roman Adler. Wolfgang Bilger and Elisabeth Kaltenegger gave valuable comments on the experimental design and abundance of alkaloids. We acknowledge funding of climate chambers by the DFG (INST 257/607-1 FUGG).

## 6 Author Contributions

KSCHR designed, coordinated and supervised the research, compiled the seed material, performed the statistical analyses and wrote the manuscript. LP, BE, MA propagated the plants, applied the experimental treatments, acquired data on plant growth and morphological functional traits and prepared the samples for metabolome analyses. TD planned and conducted the metabolomics analyses based on the FT-ICR-MS, TD and SG planned and performed the verification experiments on the UPLC-TQ-MS/MS. TD, AE and KSCHW contributed to the final version of the manuscript and all authors approved the final manuscript version.

## 7 Data availability

All data related to this article will be published in Dryad once the article is accepted for publication. Metabolome data will additionally be deposited in MetaboLights. R-codes will be published in GitHub.

## 11 Supporting Information

### SI.1: Overview of the experimental setup

SI. 1: Overview of the experimental setup
| Latitudinal Origin | Stress Treatment Combination | Population Identity | Number Maternal Families |
| --- | --- | --- | --- |
| OS | D0 - H0 | AL | 7 |
| OS | D0 - H0 | FI | 6 |
| OS | D0 - H0 | PO | 6 |
| OS | D0 - H0 | TO | 5 |
| OS | D0 - H1 | AL | 7 |
| OS | D0 - H1 | FI | 6 |
| OS | D0 - H1 | PO | 6 |
| OS | D0 - H1 | TO | 5 |
| OS | D1 - H0 | AL | 7 |
| OS | D1 - H0 | FI | 6 |
| OS | D1 - H0 | PO | 6 |
| OS | D1 - H0 | TO | 5 |
| OS | D1 - H1 | AL | 7 |
| OS | D1 - H1 | FI | 6 |
| OS | D1 - H1 | PO | 6 |
| OS | D1 - H1 | TO | 5 |
| ON | D0 - H0 | KF | 7 |
| ON | D0 - H0 | KH | 1 |
| ON | D0 - H0 | KS | 6 |
| ON | D0 - H0 | SP | 3 |
| ON | D0 - H1 | KF | 7 |
| ON | D0 - H1 | KH | 1 |
| ON | D0 - H1 | KS | 6 |
| ON | D0 - H1 | SP | 3 |
| ON | D1 - H0 | KF | 7 |
| ON | D1 - H0 | KH | 1 |
| ON | D1 - H0 | KS | 6 |
| ON | D1 - H0 | SP | 3 |
| ON | D1 - H1 | KF | 7 |
| ON | D1 - H1 | KH | 1 |
| ON | D1 - H1 | KS | 6 |
| ON | D1 - H1 | SP | 3 |

The experiment comprised eight latitudinal origin (O_N_ = north, O_S_ = south) × water treatment (D_0_ = control, D_1_= drought) × temperature treatment (H_0_ = control, H_1_ = heat) combinations with 4 replicate populations for each origin and up to 7 replicate maternal families. The number of replicate families was determined by the availability of vital seedlings, which was restricted for populations KH and SP due to low germination and high mortality rates. The experiment comprised four plants derived from the same family within a population, which were assigned to the four stress treatment combinations.

### SI.2: Overview of the most important MS parameter settings

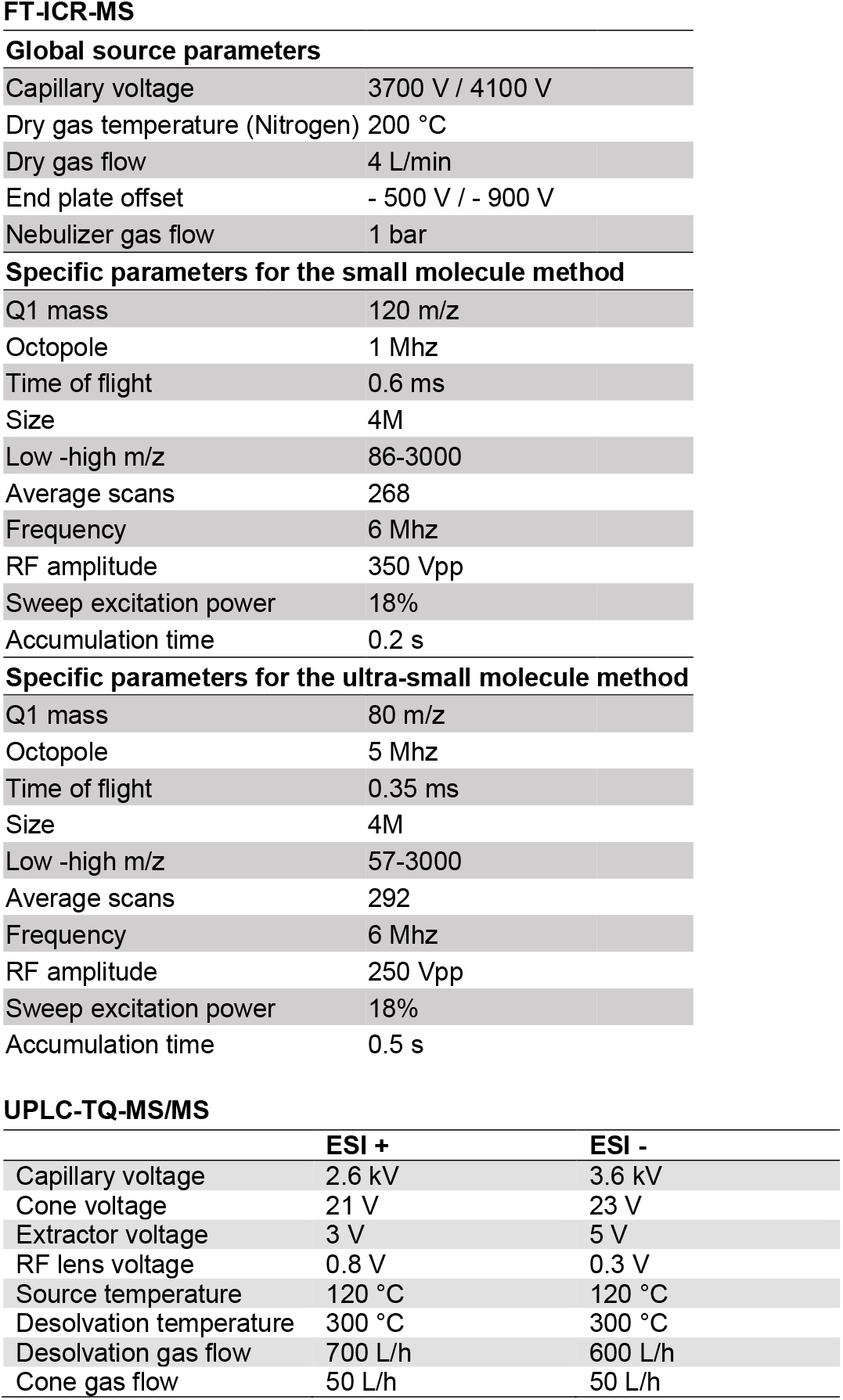

|  |  |
| --- | --- |
| Capillary voltage | 3700 V / 4100 V |
| Dry gas temperature (Nitrogen) | 200 °C |
| Dry gas flow | 4 L/min |
| End plate offset | - 500 V / - 900 V |
| Nebulizer gas flow | 1 bar |
| **Specific parameters for the small molecule method** |  |
| Q1 mass | 120 m/z |
| Octopole | 1 Mhz |
| Time of flight | 0.6 ms |
| Size | 4M |
| Low -high m/z | 86-3000 |
| Average scans | 268 |
| Frequency | 6 Mhz |
| RF amplitude | 350 Vpp |
| Sweep excitation power | 18% |
| Accumulation time | 0.2 s |
| **Specific parameters for the ultra-small molecule method** |  |
| Q1 mass | 80 m/z |
| Octopole | 5 Mhz |
| Time of flight | 0.35 ms |
| Size | 4M |
| Low -high m/z | 57-3000 |
| Average scans | 292 |
| Frequency | 6 Mhz |
| RF amplitude | 250 Vpp |
| Sweep excitation power | 18% |
| Accumulation time | 0.5 s |

**UPLC-TQ-MS/MS**
|  | **ESI +** | **ESI -** |
| --- | --- | --- |
| Capillary voltage | 2.6 kV | 3.6 kV |
| Cone voltage | 21 V | 23 V |
| Extractor voltage | 3 V | 5 V |
| RF lens voltage | 0.8 V | 0.3 V |
| Source temperature | 120 °C | 120 °C |
| Desolvation temperature | 300 °C | 300 °C |
| Desolvation gas flow | 700 L/h | 600 L/h |
| Cone gas flow | 50 L/h | 50 L/h |

### SI.3: Overview for model optimization procedure for the targeted statistical analyses of metabolome data

The Intensities of 341 pre-selected (log_2_-fold change either >=1 or <=-1) metabolites (M) were fitted as response variable in models containing the fixed effects origin (O), water treatment (D) and temperature treatment (H) and the random effect population (P). Each metabolite ran with 13 alternative models differing in i) the response data transformation (untransformed, natural logarithm=**log**, square root=**sqrt**), ii) error family (**gaussian, poisson, nbinom1** = over/underdispersed Poisson with quadratic parametrization, **nbinom2** = over/underdispersed poisson with linear parametrization), iii) dispersion formulas (**dispformula**) to account for variance inhomogeneity across treatment combination groups, and iv) zero-inflation formulas (**ziformula**) to account for an access of zeroes in the data. We extracted the results from the nonparametric dispersion test (i.e., residuals *versus* fitted), Kolmogorov-Smirnov test for uniformity, and Levene’s test for homogeneity of variance, which are provided for targeted model optimization in the R-package “DHARMa” (Hartig, 2021) for all 13 models. We regarded the simplest model with negative results for all three tests as the best fit model. Significance tests for the combined effects of origin, water treatment and temperature were exclusively done for the best-fit model.

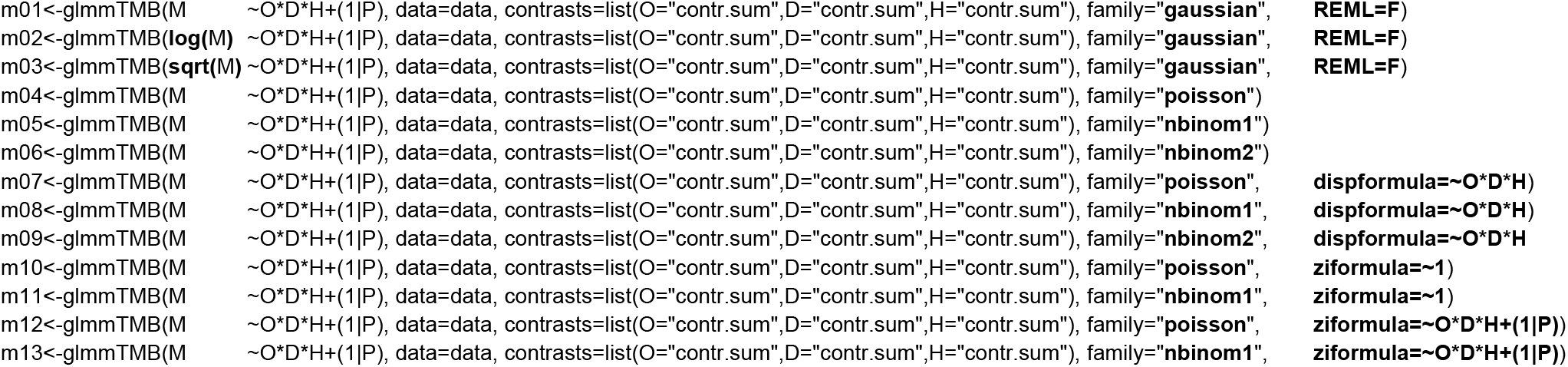

### SI.4: Results of targeted statistical metabolome analysis based on uncorrected p-values

Please see attached Excel document.

### SI.5: Evidence for the abundance of loline in *Cakile*

**Fig. SI.**
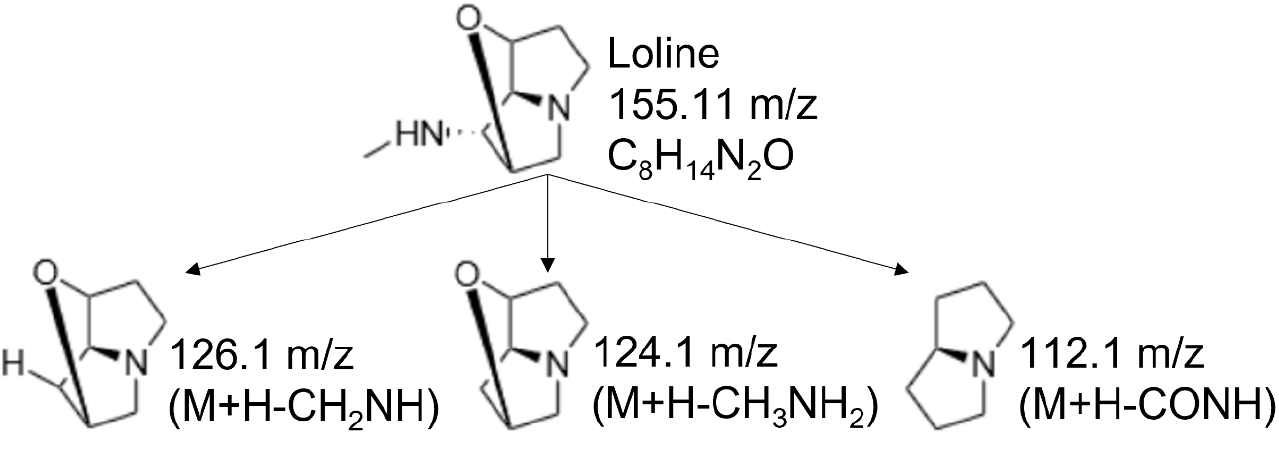
5-1: Structure and fragmentation of loline according to Adhikari (2016)

**Tab. SI.** 5-1: Predicted and observed masses, sum formulas and MRM fragments for loline.

|  | Mass | Sum formula | Fragment 1 m/z | Relative intensity fragment 1 | Fragment 2 m/z | Relative intensity fragment 2 | Fragment 3 m/z | Relative intensity fragment 3 |
| --- | --- | --- | --- | --- | --- | --- | --- | --- |
| Predicted | 155.1 | $C_8H_{14}N_2O$ | 126.1 | 2.60 | 124.1 | 9.81 | 112.1 | 22.62 |
| Observed | 155.11 | $C_8H_{14}N_2O$ | 126.09 | 2.86 | 124.07 | 12.26 | 112.07 | 69.69 |

**Tab. SI.**
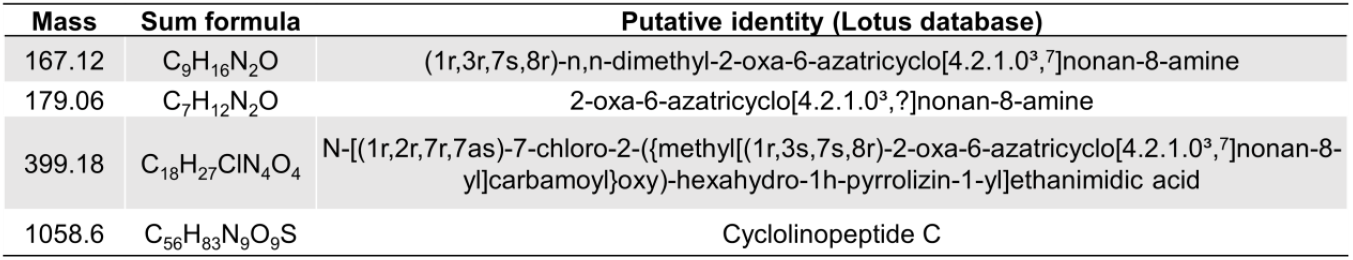
5-2: Other metabolites in leaf extracts of *Cakile*, which are putatively associated with loline alkaloid metabolism as detected with FT-ICR-MS analyses.

